# The Mediator complex subunit Med19 extends healthy lifespan in *Drosophila* by preventing cellular and organismal frailty

**DOI:** 10.1101/2025.06.18.659864

**Authors:** Adeline Payet, Emmanuelle Guillou, Sandra Bernat-Fabre, Cécile Dromard-Berthézène, Xavier Contreras, Alexia Rivero, Denis Jullien, Henri-Marc Bourbon, Jean-Louis Frendo, Coralie Sengenes, Muriel Boube

## Abstract

Aging involves a progressive decline in physiological functions, often marked by the onset of a “frailty point” just before survival rates decrease rapidly. Here, we investigate how the Mediator subunit Med19 modulates this transition in *Drosophila*. We find that upregulating *Med19* extends the median lifespan by nearly 90% and postpones the onset of accelerated mortality, suggesting that Med19 helps preserve the resilience phase of aging. In contrast, Med19 downregulation sharply reduces both median and maximum lifespan, advancing the frailty threshold. We show that Med19 knockdown increases fly vulnerability to environmental insults such as oxidative and genotoxic challenges whereas Med19 upregulation helps them resist these stresses, underscoring Med19’s protective role in maintaining genomic integrity. We link these phenotypes to altered stress-response pathways at the cellular level: Med19-depleted cells show a compensatory upregulation of genes involved in iron–sulfur cluster biogenesis, glutathione metabolism, and DNA damage repair. At the cellular level, Med19 depletion triggers a “loser” phenotype in cell competition assays, activating the JNK pathway and undergoing apoptosis, highlighting a form of “cellular frailty” that parallels organismal frailty. Finally, we found that the Med19 protein level naturally decreases with age and showed that restoring Med19 expression in aged flies increases fitness and delays the onset of frailty even in older, “frail” individuals, underscoring its significance as an aging regulator. Altogether, our findings establish Med19 as a crucial mediator of lifespan and stress resilience, suggesting it acts as a rheostat that modulates the transition from healthy aging to frailty in *Drosophila*.

**GRAPHICAL ABSTRACT:** 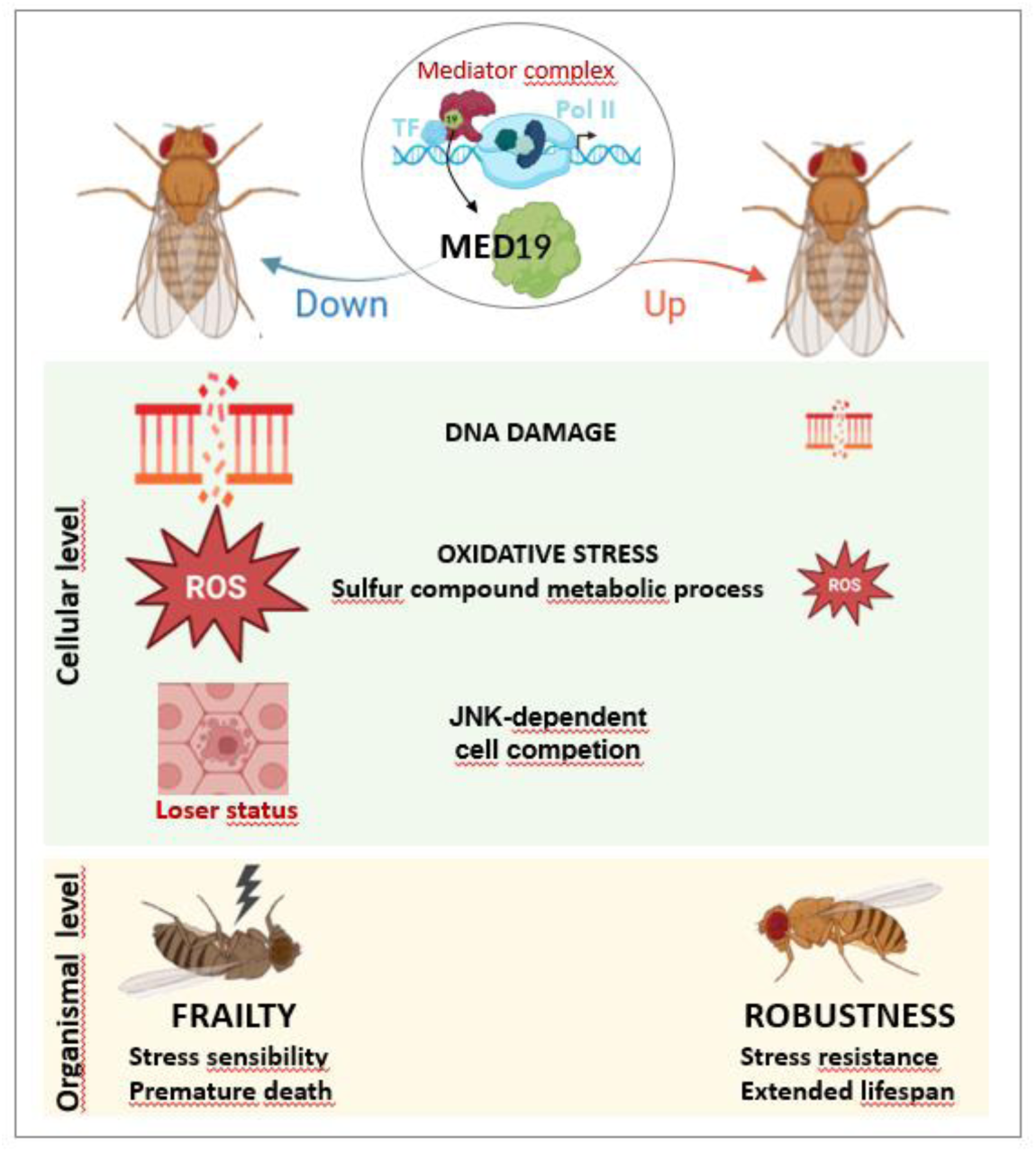

*Highlights:*

- At the organismal level, Med19 upregulation extends *Drosophila* lifespan by 90%, delays the onset of frailty and enhances oxidative and DNA damage stress resistance, while its depletion has the opposite effect.
- Med19 depletion alters the expression of stress response genes and induces a loser-cell phenotype in cell competition assays revealing its crucial function in controlling cellular fitness.
- Med19 protein levels naturally decline with age, and its restoration in aged flies improves both healthspan and lifespan.

## INTRODUCTION

Aging is a multifaceted process marked by progressive declines in cellular and physiological functions, culminating in increased susceptibility to stress and disease (López-Otín et al. 2013). In many organisms, aging is characterized by the appearance of a “frailty point,” after which mortality rises precipitously (Clegg et al. 2013). Understanding how organisms delay this transition and maintain resilience has become the next challenge in aging research, aiming to prolong not only lifespan but also healthspan. Among the hallmarks of aging, oxidative damage and genomic instability stand out as significant drivers of functional decline (López-Otín et al. 2013; Gómez et al. 2023; Lidzbarsky et al. 2018). *Drosophila* has been a particularly insightful model, illustrating that improvements in mitochondrial function and DNA repair can slow the aging process, whereas impairments in these pathways hasten frailty (Tsurumi & Li 2020; Piper & Partridge 2018; Vermeulen & Loeschcke 2007). At the cellular level, *Drosophila* also demonstrates that the active elimination of frail cells via cell competition - a mechanism conserved in humans - is crucial for maintaining tissue homeostasis and delaying age-related degeneration (Merino et al. 2016; Nagata & Igaki 2024).

Less is known, however, about the transcriptional regulators that integrate these processes to ensure robust organismal survival over time. The Mediator complex (MED)—a 25–30-subunit assembly conserved from yeast to humans—links DNA-bound transcription factors (TFs) to RNA polymerase II (RNAPII) integrating various aspects of gene regulation within the nuclear compartment. MED influences transcription initiation, elongation, and chromatin organization (Richter et al. 2022; Boube et al. 2002). Despite the broad role of MED complex in regulating most RNAPII-driven genes, growing evidence suggests that specific subunits exert specialized functions in disease and development (Schiano et al. 2014; Ilchuk et al. 2023). The Med19 subunit, in particular, is significantly upregulated in multiple cancers—among them lung, liver, and hepatocellular carcinoma—and is linked to tumor growth and metastasis (Zhang et al. 2021; Jin et al. 2024). However, the molecular bases of Med19’s tissue- and context-specific actions remain largely uncharacterized. Previous work in *Drosophila* established that Med19 physically interacts with developmental regulators such as HOX and GATA TFs, influencing their target gene expression *in vivo* (Boube et al. 2014; Immarigeon et al. 2020; Jullien et al. 2022). Yet, the potential roles of Med19 in adult physiology, stress resistance, and lifespan regulation have been poorly explored.

Here, by combining genetic, transcriptomic, and functional assays, we establish that Med19 modulates stress-response pathways, influencing both organismal survival and cellular fitness. At the organism level, Med19 upregulation greatly extends lifespan, postpones the onset of frailty and enhances oxidative and DNA damage tolerance. By contrast, its depletion reduces lifespan and displays higher susceptibility to environmental challenges. At the cellular level, Med19 depletion alters expression of stress-response genes involved in DNA damage response, iron–sulfur cluster biogenesis and glutathione metabolism. It also triggers a loser-cell phenotype in cell competition assays—an indicator of “cellular frailty”. Importantly, we found that Med19 protein levels naturally decline in aged flies, and that late induction of Med19 in already aged individuals further prolongs survival and preserves fitness. Our study provides new insights into how a Mediator complex subunit can fine-tune fundamental processes that underlie healthy aging.

## RESULTS

### Med19 is a critical regulator of *Drosophila* lifespan delaying the onset of frailty

We previously showed that ubiquitous, expression of Med19 (driven at low level by an *ubMed19* transgene) partially rescued the lethality of *Med19* knockout (KO) individuals (Boube et al. 2014). Interestingly, we further noticed that otherwise wild-type flies bearing the *ubMed19* transgene exhibit a striking extension in lifespan. To measure this effect, we performed survival analysis and observed a drastic increase in the median lifespan of *ubMed19* individuals by almost 90% (p < 0.0001) and in the maximum lifespan by 51% (Figure 1A-A’ and S1 table). The maximum lifespan reflects the age reached by the longest-lived individuals, while the median lifespan refers to the age at which 50% of the population has died. The concurrent extension of both parameters suggests that Med19 overexpression not only enhances the overall physiological limits of longevity but also delays the onset of age-related decline in the general population. Analysis of the survival curves further showed no significant difference in mortality rate after the onset of death events, as indicated by comparable maximum slopes between conditions. Nevertheless, the onset of frailty i.e., defined as the transition point from a phase of relative resilience to a rapid increase in mortality risk, was dramatically delayed in *ubMed19* individuals (Figure 1A’).

**Figure 1.**
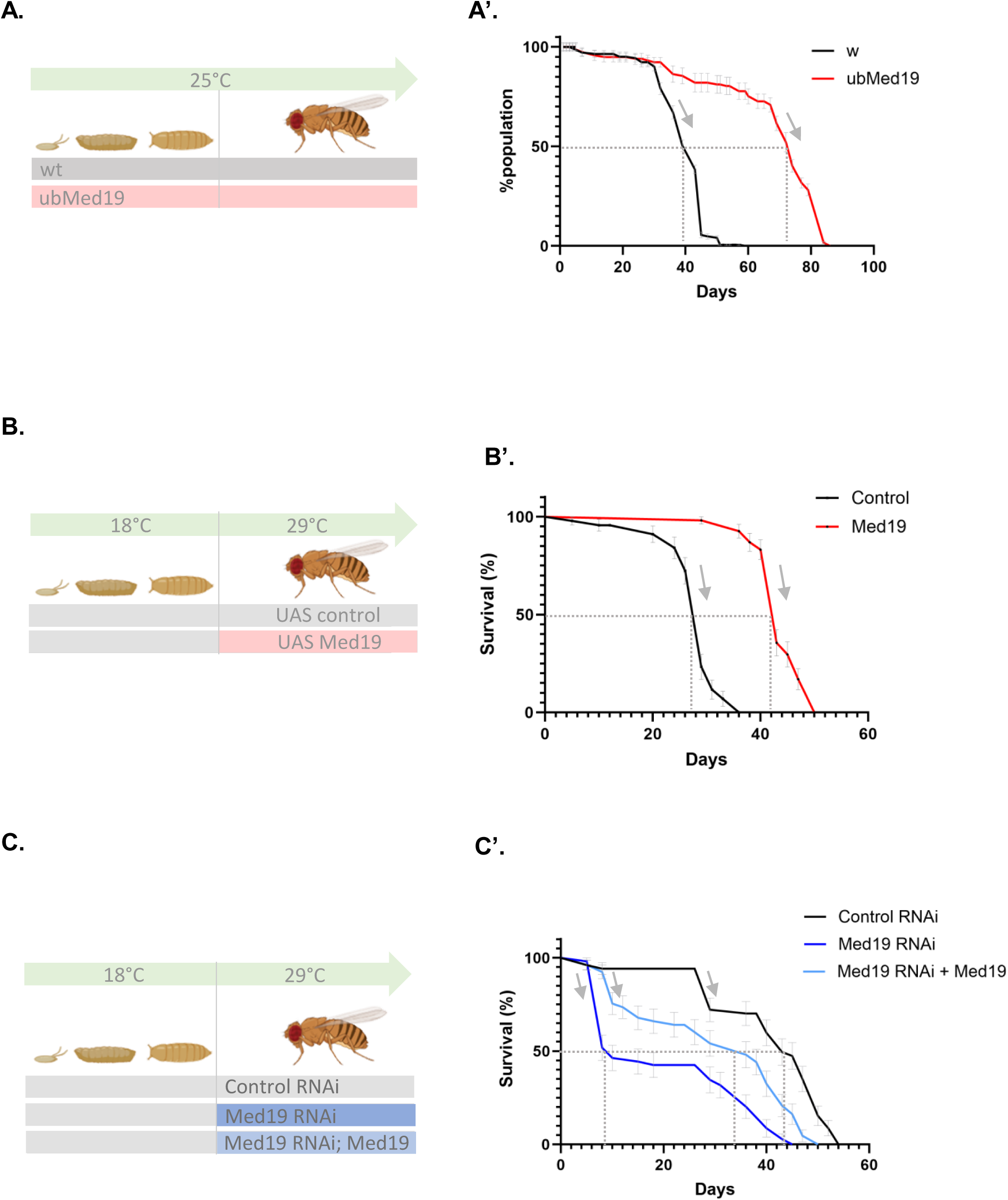
Med19 levels modulate longevity by postponing the onset of frailty. (A). Schematic representation of the constitutive upregulation of Med19 using the *ubiquitin* promoter (*ub-Med19*). (A’) Survival curve showing that flies overexpressing Med19 (Ub-Med19, red) exhibit significantly extended lifespan compared to control flies (black). Dotted lines indicate median lifespan and arrows, the curve slopes after curve inflection points. See also S1 table for survival curve statistical analysis. (B) Inducible overexpression of Med19 protein using the UAS/Gal4 system. Flies were raised at 18°C to suppress Gal4 activity by the GAL80ts, then shifted to 29°C to induce Med19 expression in adults. (B’) Survival analysis reveals that adult-induced overexpression of Med19 (red) extends median and maximal lifespan and displace the inflection point of survival curves (arrows) relative to *UAS-Control* flies (black), confirming that Med19 positively regulates longevity when activated during adulthood. See also S1 table. (C) RNAi-mediated knockdown of Med19 using the UAS/Gal4 system with adult-specific induction (Gal80ts). Flies were raised at 18°C and shifted to 29°C to activate Med19 RNAi expression. (C’) Lifespan analysis shows that Med19 knockdown (dark blue) significantly reduces median (dotted lines) and maximal lifespan compared to control RNAi (black) and reduces the curve inflection point meaning that the transition to rapid mortality acceleration (arrows) occurs earlier in the Med19 KD condition indicating a deleterious effect on longevity. Co-expression of Med19 protein with *Med19* RNAi (light blue) partially rescues the lifespan reduction, confirming Med19 specific involvement in regulating lifespan. See S1 table.

To assess whether Med19 overexpression during adulthood is sufficient to extend lifespan, we used the temperature-sensitive UAS / GAL4 / GAL80ts system to induce ubiquitous Med19 upregulation specifically in adult flies, an induction previously shown to rescue cellular lethality of *Med19* mutant clones (Boube et al. 2014). As shown in Figure 1B-B’ and S1 table, adult-specific upregulation of Med19 led to a significant increase in median lifespan by 48% (p < 0.0001) and in maximum lifespan by 51% (p < 0.01). As observed in *ubMed19* individuals, adult-specific Med19 expression delayed the onset of accelerated mortality without altering the mortality rate once engaged (Figure 1B-B’ and S1 table). These results support a protective role for Med19 in aging by delaying the transition to frailty even when expression is activated during adulthood.

We then tested whether Med19 knockdown (KD, using interfering RNA) also impacted lifespan. Adult-specific Med19 downregulation severely decreased median and maximum lifespan by nearly 70% (p < 0.0001) and 14% (p < 0.05) respectively compared to control flies. Conversely to Med19 overexpression, Med19 knockdown shifted the onset of frailty markedly earlier than in controls, as indicated by an earlier inflection point in the survival curve (Figure 1C-C’ and S1 table). Partial rescue by Med19 co-expression confirmed Med19-specific involvement in regulating lifespan and frailty onset.

Taken together, our data establish Med19 as a key modulator of longevity, acting as a rheostat for the onset of frailty.

### At the organismal level, Med19 influences resistance to stress

A key feature of frailty is diminished resilience to stressors, resulting in higher vulnerability to adverse health outcomes. We therefore sought to investigate the role of Med19 in regulating stress resistance at the organismal level. To do so, we exposed adult individuals to two stress paradigms: (1) short pulses of UV irradiation (UV) which creates lesions such as pyrimidine dimers that disrupt DNA integrity and (2) exposure to paraquat (PQ), a well-characterized herbicide that enhances mitochondrial superoxide production and stimulates manganese-superoxide dismutase activity, leading to increased cellular ROS levels. We then assessed the effects of Med19 knock-down or overexpression under these conditions.

Upon UV irradiation, median lifespan is reduced by ∼20% (compare Figure1C’ to Figure 2A’). In the same conditions, RNAi-mediated Med19 knockdown in adults drastically reduced median and maximal lifespan by 67% (p < 0.0001) and 27% (p < 0.001), respectively (Figure 2A-A’ and S1 table). This reduction was partially rescued by restoring Med19 expression confirming the specific role of Med19 in protecting against DNA damage-induced stress. These results highlight Med19 as a key factor in genotoxic insults resistance.

**Figure 2.**
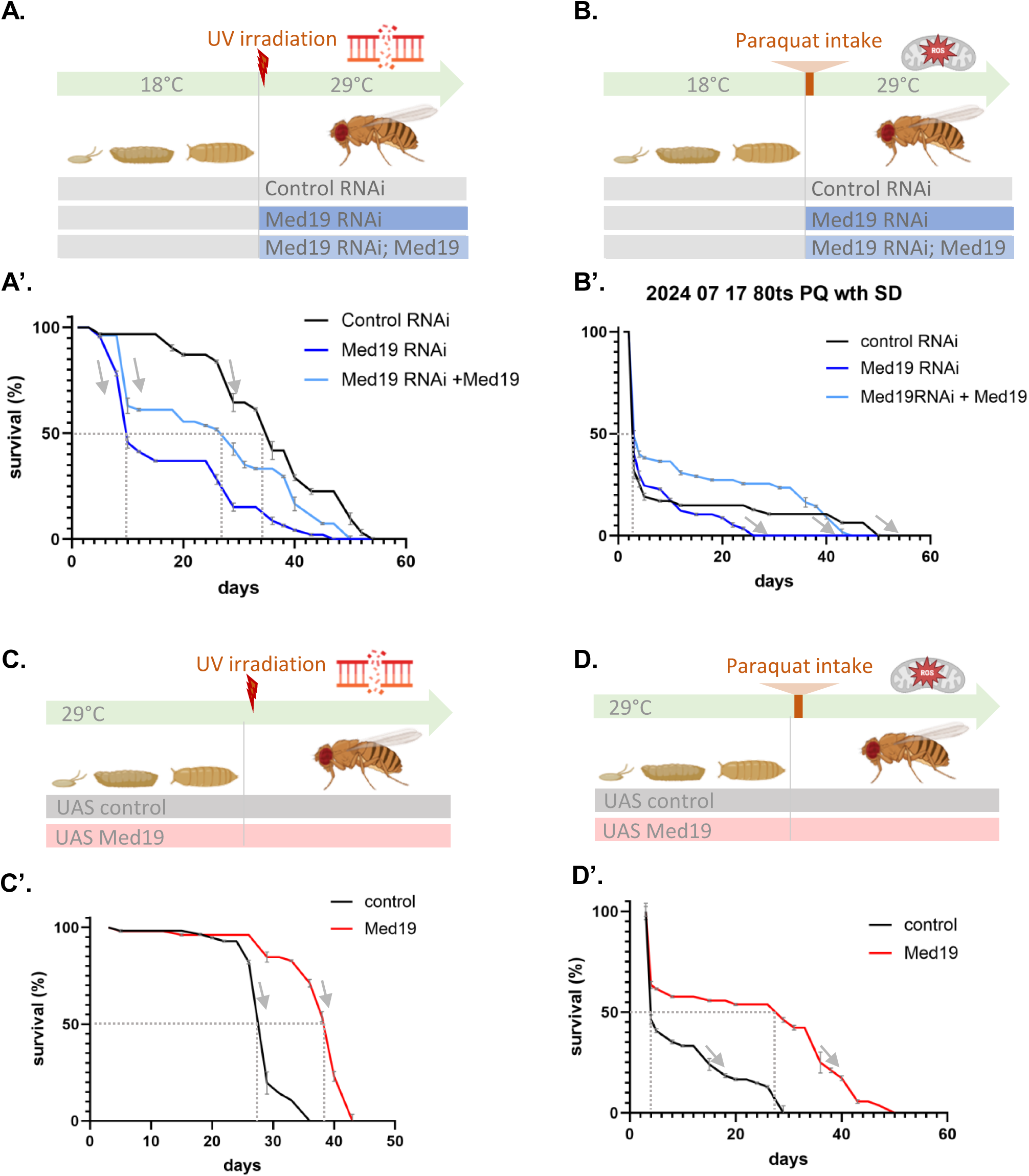
Med19 level modulates resistance to DNA damage and oxidative stress. (A) Schematic of the experimental design for Med19 knockdown using RNAi, with adult-specific induction at 29°C after exposure to UV irradiation. (A’) Survival curves demonstrate that flies with *Med19* RNAi (blue) exhibit reduced survival under UV-induced stress compared to control RNAi (black). Dotted lines indicate median lifespan and arrows, curve slopes after curve inflection points. See also S1 table for survival curve statistical analysis. Co-expression of Med19 with RNAi (light blue) partially rescues this sensitivity confirming the specificity of Med19 in UV stress resistance. See S1 table for survival statistics. (B) Experimental setup for Med19 knockdown with exposure to 20mM paraquat, a chemical that induces oxidative stress. (B’) Corresponding survival analysis for flies expressing Med19 RNAi (dark blue), Med19 RNAi + Med19 upregulation (light blue) or control RNAi (black). Dotted lines indicate median lifespan and arrows, curve slopes after curve inflection points. See also S1 table for survival curve statistical analysis, respectively. These results indicate a Med19’s role in oxidative stress tolerance. See S1 table for survival statistics. (C) Schematic of the experimental design for continuous Med19 upregulation with a *ubGAL4 UAS-Med19* transgene combination (red), and UV exposure at the beginning of adult stage. (C’) Survival analysis reveals that Med19 upregulating flies (red) show markedly enhanced survival compared to control flies (black) when subjected to UV irradiation, indicating that Med19 upregulation confers protection against DNA damage. Dotted lines indicate median lifespan and arrows, changes in curve inflection point reflecting the onset of frailty. See S1 table for survival statistics. (D) Experimental design for Med19 overexpression and subsequent exposure to 10mM paraquat.(D’) Flies overexpressing Med19 (red) also show significantly improved median lifespan (35 vs 24 days for control) under oxidative stress induced by paraquat compared to controls (black), indicating that increased Med19 levels enhance resistance oxidative stress. See S1 table for survival statistics.

Massive lethality occurred shortly after paraquat exposure both in control and Med19 RNAi flies, resulting in no significant differences in median or maximal lifespan between conditions. However, the mean lifespan of Med19 RNAi flies was significantly reduced by 48% compared to control RNAi (p < 0.05; Figure 2B-B’ and S1 Table). Mean lifespan was restored to control levels upon introduction of a Med19-expressing transgene, further establishing the key role of Med19 in mitigating the impact of oxidative stress on longevity.

In contrast, Med19 upregulation conferred a marked protective effect. Upon UV stress, Med19 overexpression improved mean lifespan by 38% (p < 0.0001) with a clear delay in the curve inflection point (Figure 2C-C’ and S1 Table). Under paraquat-induced oxidative stress, although initial lethality was similar between groups, survivors upregulating Med19 displayed a delay in the onset of accelerated mortality (Figure 2D-D’ and S1 Table). These flies showed an 86% extension of median lifespan (p < 0.0001; Figure 2D-D’ and S1 Table). These findings indicate that higher Med19 levels bolster defense mechanisms against DNA damage and oxidative stress.

Together, these results indicate that Med19 strengthens resilience against genotoxic and oxidative stress, reinforcing its role in maintaining stress resistance during aging and modulating lifespan and frailty trajectories.

### Med19: a key transcriptional regulator of cellular stress response

To better understand Med19 role in modulating frailty, we searched to identifying the set of genes regulated by *Med19* at the transcriptional level. To this end, we used RNA-seq data performed on larval wing discs expressing either a Med19-specific RNAi or a control Luciferase (Luc) RNAi together with Dicer2 (Dcr2) promoting RNAi efficiency (Figure 3A). Bioinformatic analysis identified 786 differentially expressed (DE) genes, based on their expression level (Log2 fold change >0,5 and <-0,5) and their significance (adjusted p-value <0,05) (Figure 2B). Of these, 276 genes were down-regulated and 510 upregulated (Figure 3B). These results confirm a specific role of this Mediator complex subunit in differential regulation, negative or positive, of small sets of genes, following our analysis of Med19 target genes using an auxin induced Med19 degradation (Jullien et al. 2022). While we already shown that many downregulated targets were associated with the transcriptional control of development (Jullien et al. 2022), we focused here on the upregulated genes, which revealed links to stress response networks. Gene ontology (GO) analysis of the upregulated genes showed that Med19 depletion results into overexpression of several genes falling into the GO terms GO:0006974 “DNA damage response” and GO:0006780 “sulfur compound metabolic processes” (Figure 2C).

**Figure 3.**
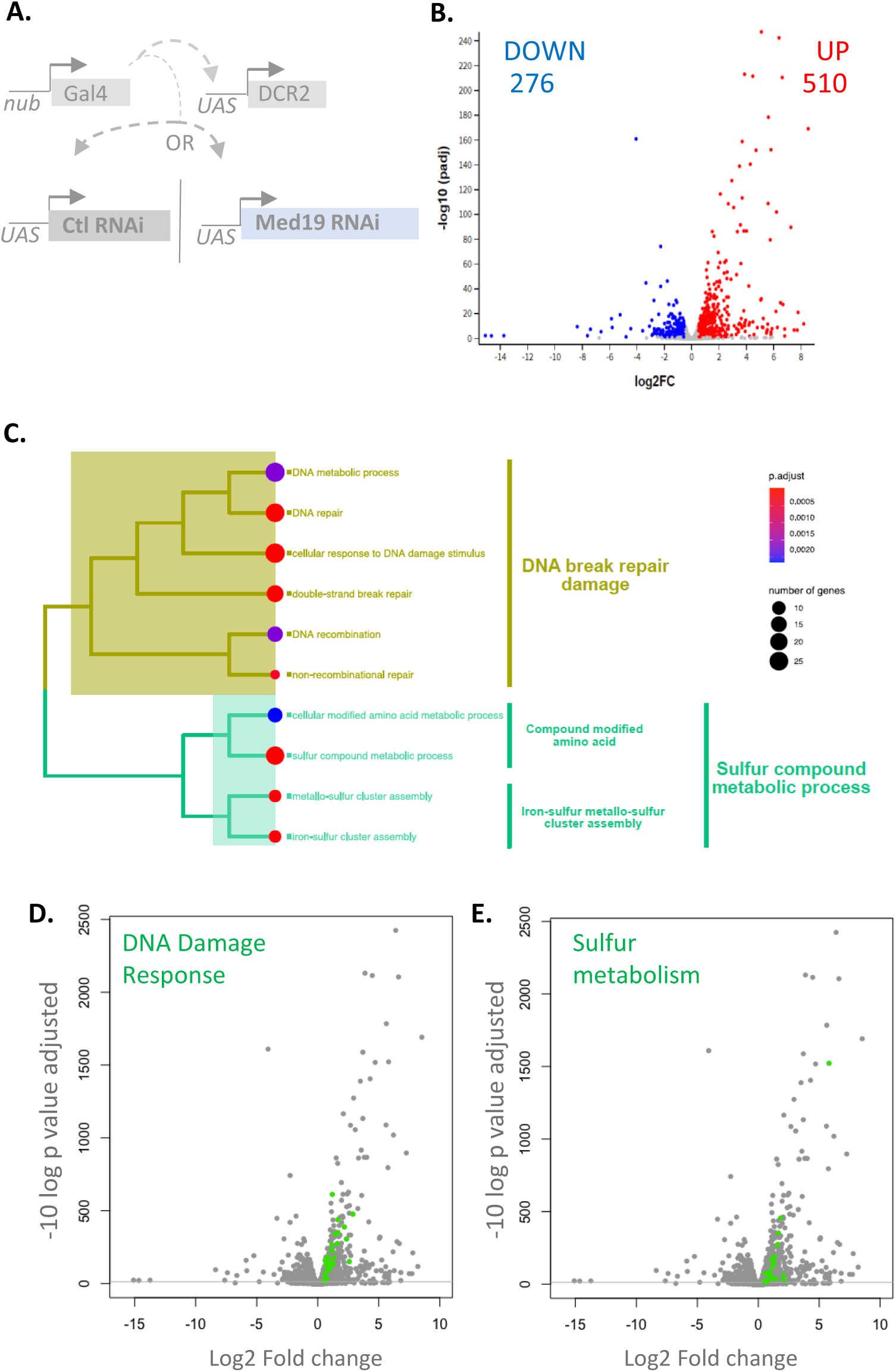
Identification of genes regulated by Med19. (A) Experimental design for transcriptomic analysis. Either a control UAS RNAi line (against Luciferase) or a *UAS-Med19* RNAi together with *UAS-Dicer2* (DCR2) were expressed under the control of a strong *nubGAL4* inducer in wing imaginal pouch to assess the effect of Med19 knockdown on gene expression profiles. (B) Volcano plot of differentially expressed genes in larvae wing pouch expressing Med19 RNAi or Luc RNAi. Genes are highlighted: in red if their log2 Fold Change is > 0.5 and their adjusted p-value is <0.05, in blue if their log2 Fold Change is < -0.5 and their adjusted p-value is <0.05. (C) Gene Ontology (GO) enrichment analysis performed exclusively on the 510 upregulated genes reveals significant enrichment in pathways related to DNA break repair/damage response and sulfur compound metabolic processes. The heatmap color gradient indicates the significance level of enrichment, while circle size reflects the number of genes involved in each category. (D-F) Volcano plots highlighting specific upregulated stress-response pathways: (D) GO DNA Damage Response (E) GO sulfur compound metabolic processes

Genes belonging to the GO “DNA damage response” include genes activated in the context of genotoxic stress, particularly in response to double strand breaks (Figure 3D and S2 Table). Their upregulation suggests that Med19 depletion leads to altered genomic integrity in cells consistent with the reduced resistance observed at the organismal level.

Genes found in the GO “Sulfur compound metabolic process” are involved in Iron sulfur (Fe-S) cluster assembly or in glutathione metabolic process (Figure 3E and S2 Table). Fe-S clusters are essential cofactors of numerous enzymes, notably those involved in cellular respiration within the mitochondrial respiratory chain and some key regulators of DNA repair (Maio N, Rouault 2020). The upregulation of genes involved in Fe-S cluster assembly may reflect a compensatory response to cluster dysfunction, potentially triggered by increased oxidative stress following Med19 downregulation. In parallel, the upregulation of genes involved in glutathione metabolism—central to cellular antioxidant defense—further supports the notion of redox imbalance. Together, these changes suggest that Med19 depletion disrupts cellular homeostasis and triggers transcriptional adaptations aimed at maintaining mitochondrial function and countering oxidative stress.

Taken together our data reveal that Med19 depletion significantly impacts several cellular processes linked to stress response. The upregulation of genes involved in Fe-S cluster assembly and glutathione metabolism suggests that Med19 depletion may lead to oxidative stress and iron metabolism dysfunction. Furthermore, the activation of DNA damage responses genes points to a role of Med19 in maintaining genomic integrity. These results suggest a critical role for Med19 in maintaining cellular fitness through the regulation of stress responses, thereby supporting its function as a stress protector at the organismal level.

### At the cellular level, Med19-depleted cells behave as loser cells and undergo cell competition–induced apoptosis

A dedicated way to evaluate cellular frailty is cell competition, a conserved biological process in which suboptimal less-fit cells—referred to as ‘losers’—are actively eliminated by their healthier more ‘fit’ neighbors (winners) (S3 Figure A). This phenomenon plays a crucial role in tissue homeostasis, development, and aging (Merino et al. 2016). To assess whether Med19-depleted cells exhibit loser-like behavior, we investigated their ability to compete with wild-type neighbors. Our previous work highlighted a specific behavior for *Med19* knockout cells (Boube et al. 2014). We have shown that *Med19* KO clones are lethal (Figure 4A”) but survive and develop to some extent (Figure 4B”) when surrounded by cells mutant for *Minute* – affecting ribosomal proteins and displaying elevated proteotoxic stress (Baumgartner et al. 2021). This context-dependent viability of *Med19* KO cells depending on the tissue context suggested a role in cell competition.

**Figure 4:**
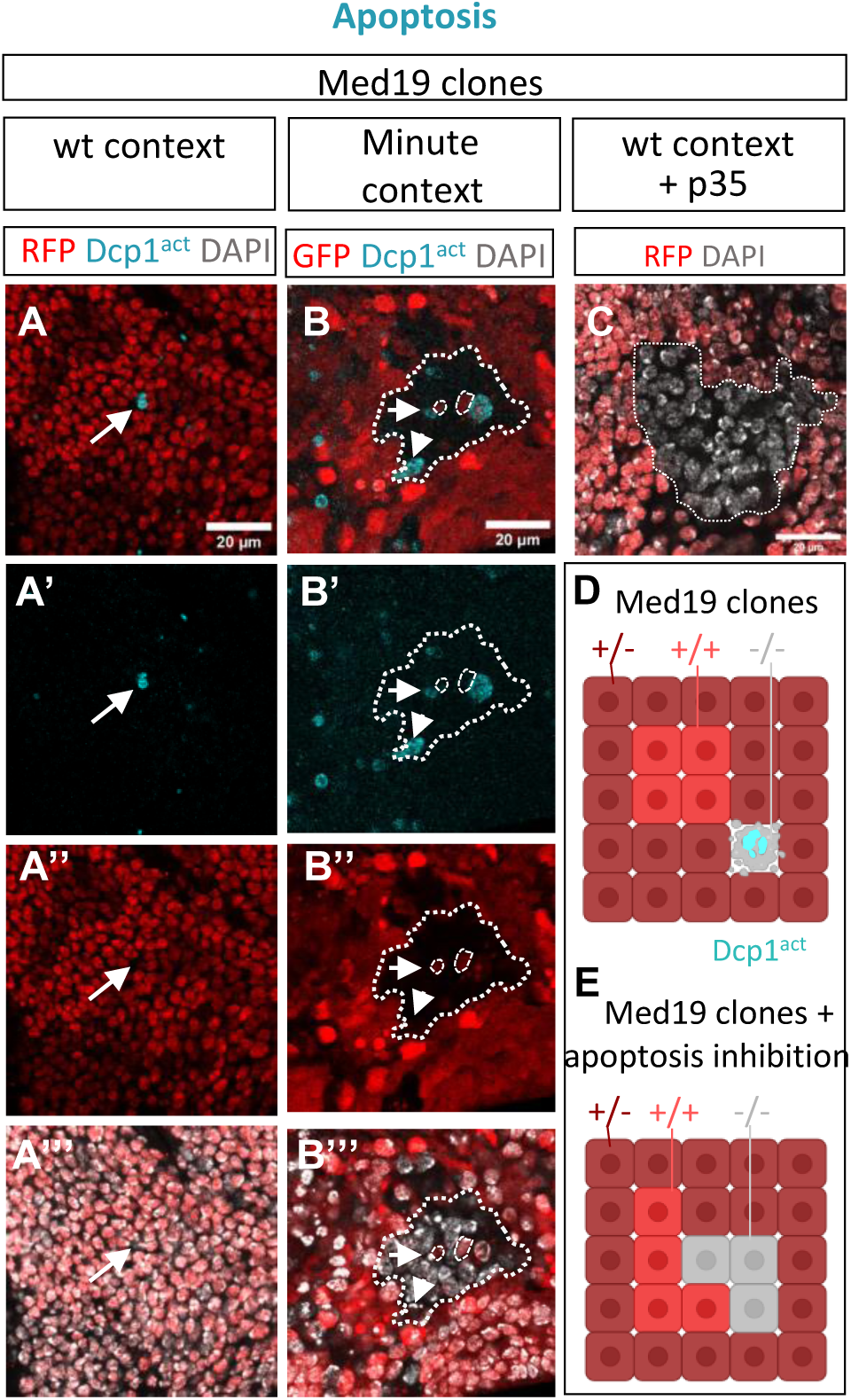
Med19-depleted cells die by apoptosis. (A-A’’’) *Med19* mutant cells are eliminated by apoptosis. (A) *Med19* mutant clones (marked by the absence of RFP) generated in a wild-type context, with immunostaining for activated caspase Dcp1 shown in blue. Single-channel Dcp1 and RFP staining are displayed in panels A’ and A’’ and merged RFP-DAPI staining in panel A’’’. Arrows indicate scattered Dcp1-positive *Med19-/-* cells adjacent to *Med19+/+* cells. (B-B’’’) *Med19*-/- clones generated in a *Minute* context. Red cells bear *Minute* mutation. *Med19-/-* clones, identifiable by the absence of GFP and presence of DAPI nuclear staining (grey) are outlined. In *Med19* mutant clones, activated Dcp1 (blue, white arrow) is observed, typically in cells situated at clone border i.e., in contact with *Med19+/+ Minute* cells (red). Single Dcp1 and GFP staining are shown in panels B’ and B’’ and double GFP-DAPI staining in panel B’’’. (C) Rescue experiments of *Med19-/-* clone lethality by blocking apoptosis. *Med19* mutant clones were generated in a wild-type context under overexpression of the caspase inhibitor P35. *Med19-/-* mutant clones (RFP-negative, DAPI-positive grey cells) are now viable. (D-E) Schematic representation of *Med19* mutant clone behavior in wild type context (D) or after apoptosis inhibition (E).

To test this hypothesis, we used the MARCM system to generate mosaic tissues containing GFP-labeled clones with mild Med19 knockdown, alongside unlabeled wild-type sibling clones (S3 Figure B). As shown in S3 Figure D-D’, Med19 knockdown clones were smaller than their corresponding wild type clones. Conversely, clones expressing a control RNAi displayed comparable size respective to their sibling wild type clone (S3 Figure C-C’). Quantitative analysis confirmed a significant reduction in Med19-depleted clone size compared to controls (p < 0.01 – S3 Figure E), supporting the idea that these cells are actively outcompeted. Importantly, when the same mild Med19 knock-down was applied to the entire tissue, wing disc size remained unaffected (S3 Figure F–I). This contrasts with the results obtained in mosaic tissue and supports the hypothesis that Med19-depleted cells are eliminated through a cell competition mechanism, rather than as a consequence of proliferation defects or intrinsic cell lethality.

Different mechanisms have been described for elimination of suboptimal “loser” cells during cell competition, such as apoptosis, engulfment by neighboring winner cells or phagocytes, extrusion from the tissue. We first investigated the hypothesis that *Med19-/-* cells die through apoptosis. To test this, we analyzed the activation of the Drosophila effector caspase Dcp1 using an antibody specific for its active cleaved form (Dcp1^act^). In our previous analysis, homozygous *Med19* KO clones were not detected (Boube et al. 2014), using activated Dcp1 staining, we were able to detect isolated apoptotic cells lacking RFP expression (Figure 4A-A’’’ and 4D). These data indicate that such *Med19* mutant cells fail to proliferate in a wild type context and are quickly eliminated from the tissue through apoptosis. In a context mutant for *Minute*, conferring a suboptimal, less fit phenotype, larger *Med19* mutant clones were generated and clear Dcp1^act^ staining was observed (Figure 4B-B”’’’). Notably, Dcp1^act^ signal was restricted to scattered cells, predominantly located at the borders of clones adjacent to *Med19^+/+^*cells, consistent with classical cell competition-induced apoptosis (arrows in Figure 4B). Supporting this, we often observed fragmented DNA in dying Dcp1^act^ positive cells (Figure 4B’’’).

To confirm that apoptosis underlies the lethality of *Med19* KO cells, we expressed the baculovirus caspase inhibitor P35 in *Med19* clones. As seen in Figure 4C and 4E, large *Med19* mutant clones are visible in the wild type context, indicating that blocking apoptosis clearly rescue *Med19* mutant cell lethality.

Taken together, our data indicate that *Med19* mutant cells are eliminated by apoptosis, especially at clone borders as expected for the elimination of loser cells during a cell competition-like mechanism.

### The JNK pathway and the cell competition marker Azot drive the elimination of Med19-depleted loser cells

Although the mechanisms by which winner and loser cells are distinguished are not fully understood, several signaling pathways have been involved. For instance, Jun N-terminal Kinase (JNK) pathway is often upregulated in loser cells and can promote their elimination by apoptosis in several cases of cell competition (Kucinski et al. 2017; Kodra et al. 2024; Igaki 2009). We tested the activation of the JNK pathway by using an antibody against the active phosphorylated form of the JNK kinase (^P^Jun). In a wild type context, all *Med19-/-* cells observed were positive to ^P^Jun staining (Figure 5A-A’’’ and 5D). In the same way, when larger *Med19-/-* clones were generated in a *Minute* context, all *Med19* mutant cells were positive for ^P^Jun signal (Figure 5B-B’’’). To determine whether JNK pathway induction is responsible for *Med19-/-* loser cell elimination, we blocked the JNK pathway using a dominant negative form of the Basket JNK kinase (BskDN). This allows the generation of large *Med19-/-* clones in wild type context, showing that JNK pathway activation is necessary for their elimination (Figure 5C and 5E).

**Figure 5.**
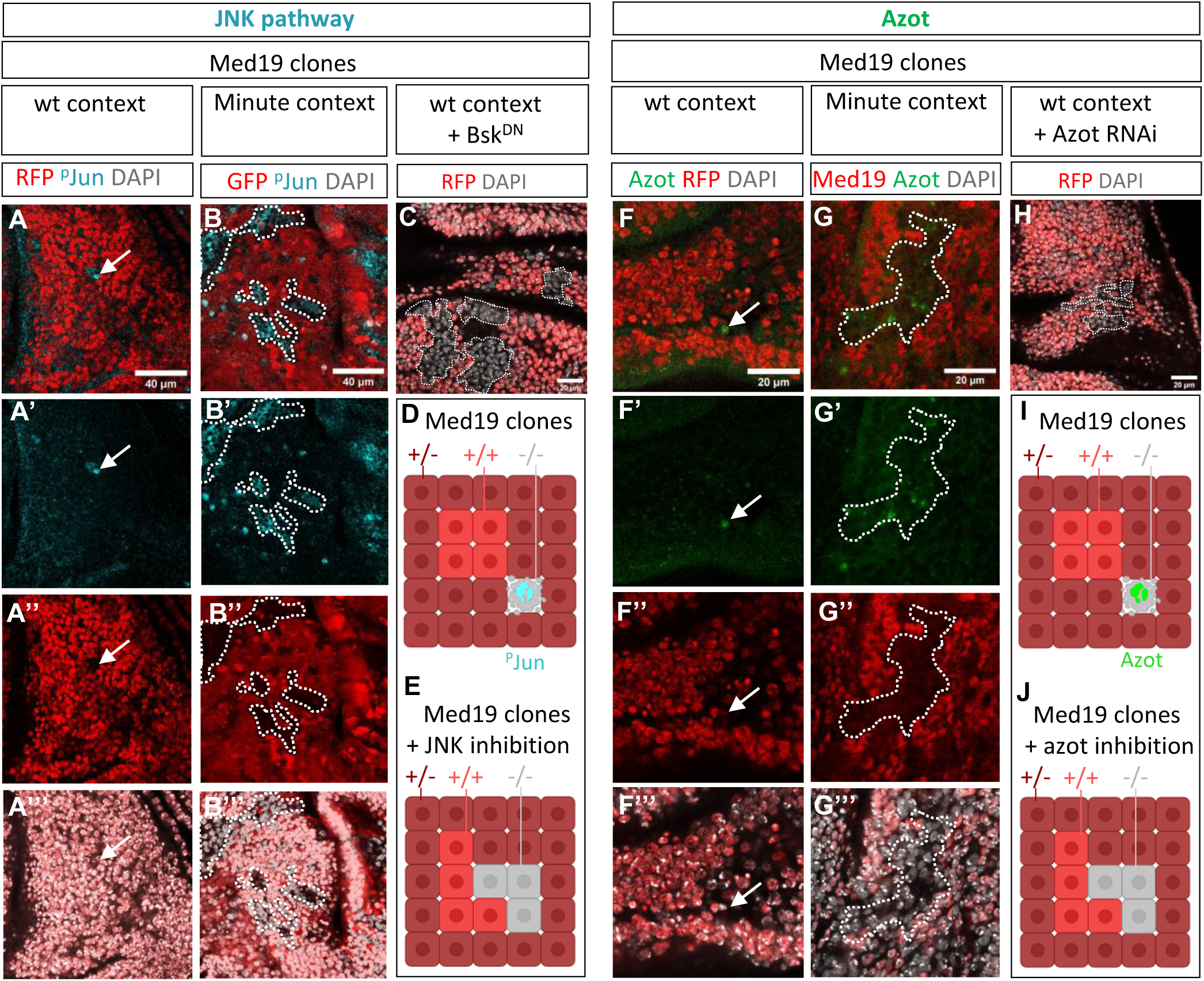
Med19-depleted cells activate the JNK pathway and the competition marker Azot. (A-A’’’) Immunostaining against phospho-Jun (^P^JUN) in *Med19-/-* clones (marked by the absence of RFP) generated in wild-type context. Arrows point to *Med19-/- cells* positive for ^P^JUN staining (cyan), indicating activation of the JNK signaling pathway. Single-channel ^P^Jun and RFP staining are displayed in panels A’ and A’’ and merged RFP-DAPI staining in panel A’’’ (B-B’’’) In a *Minute* context, immunostaining of ^P^JUN, reveals widespread activation of the JNK pathway in *Med19* mutant clones (outlined by dotted lines). (C) Rescue experiments of *Med19-/-* clone lethality by blocking JNK signaling using a dominant-negative form of the JNK Basket (Bsk), revealing the emergence of large *Med19* mutant clones (RFP-negative, DAPI-positive cells). (D-E) Schematic representation of *Med19* mutant clone behavior in wild type context (D) or after JNK pathway inhibition (E). (F-F’’’) *azot* reporter expression (green) is upregulated in *Med19-/-* clones (RFP-negative cells) generated in a wild-type context, indicating activation of *azot*, a key regulator of cell competition. (G-G’’’) Expression of the *azot-GFP* reporter (green) in *Med19-/-* clones (negative for Med19 staining-red) generated in a *Minute* context (RFP-positive cells), showing discrete induction of *azot* within *Med19* mutant clones. (H) Rescue experiments of *Med19-/-* clone lethality under RNAi-mediated *azot* inhibition. *Med19-/-* mutant clones (RFP-negative, DAPI-positive grey cells) are now viable. (I-J) Schematic representation of *Med19* mutant clone behavior in wild type context (I) or after *azot* inhibition (J).

A key actor of cell competition, the azot gene, is considered as a cell fitness checkpoint integrating information from fitness sensors such as Flower (Fwe) and Sparc proteins (Merino et al. 2015). The Azot protein is specifically expressed in loser cell and is necessary for their elimination in numerous cases of cell competition, notably those induced through *Minute* mutations and through inactivation of Dpp, JAK-STAT and Wg pathways. Using an *azotKO::GFP* reporter line, we found that *azot* is expressed in a discrete and punctiform manner in subgroups of *Med19* mutant cells in both in wild-type (Figure 5F-F’’’ and 5I) and *Minute* conditions (Figure 5G-G’’’). The use of an RNAi, mediated *azot* knockout, rescued the lethality of *Med19* KO clones in a wild-type background, further supporting its role in their elimination (Figure 5H and 5J).

Taken together, we have shown here that *Med19* KO cells are eliminated through a cell competition mechanism depending on *azot* expression, JNK activation, and apoptosis. In conclusion, Med19 induces a state of cellular frailty highlighting *Med19* as a critical regulator of cell fitness.

### Age-dependent decline in Med19 protein levels and its impact on *Drosophila* fitness

Given Med19’s role in promoting cellular and organismal robustness, we investigated whether its expression declines with age. To this end, we performed western-blot analysis comparing young and very old flies (i.e., 5 to 45 days-old at 29°C). As shown in Figure 6A-A’, we clearly observed lower amounts of Med19 protein in old flies compared to young ones.

**Figure. 6.**
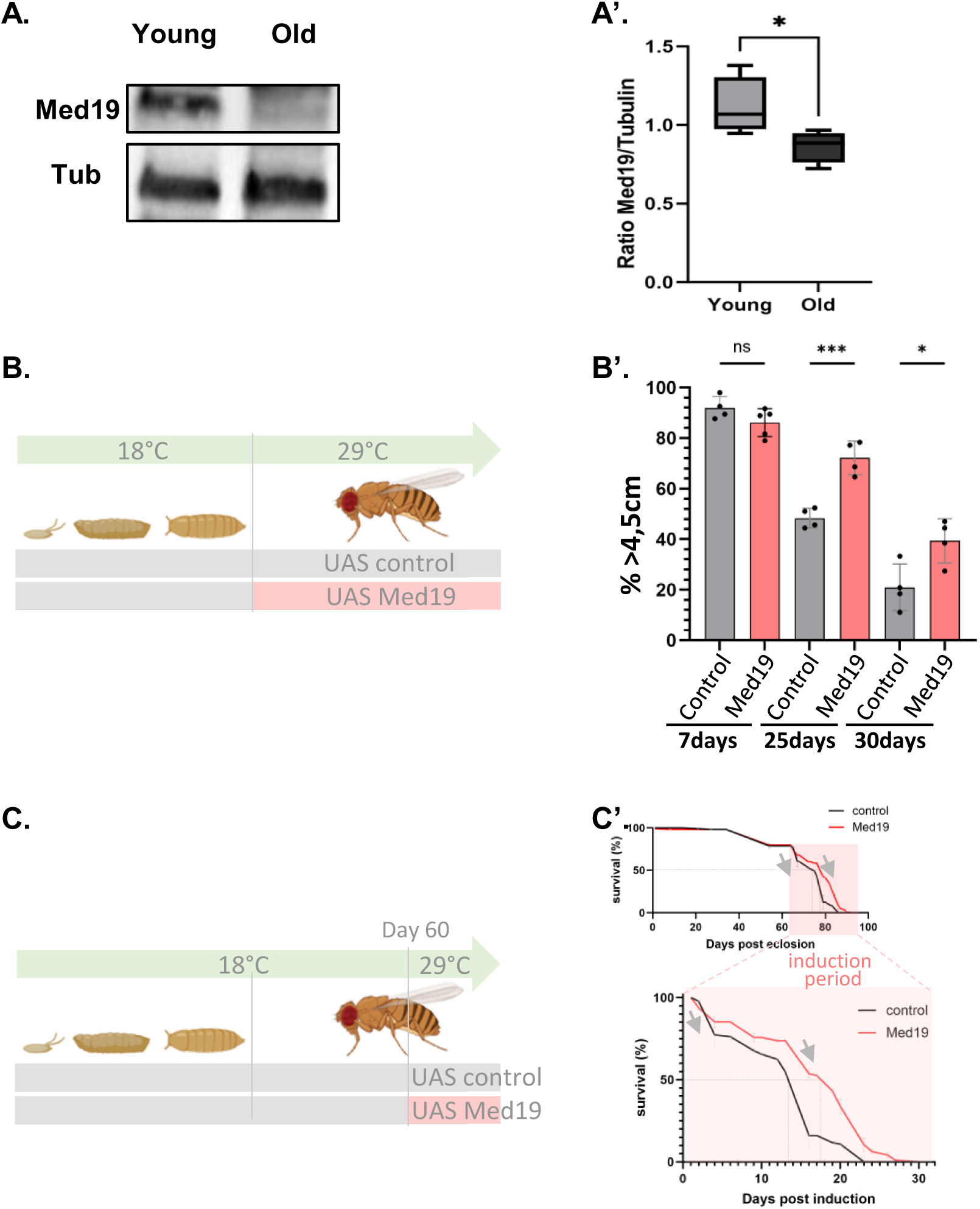
Age-dependent decline in Med19 protein levels and impacts of restoring Med19 in frail individuals. (A-A’) Med19 decreases with age in flies. Western blot and quantification of Med19 protein in young (3-days-old) and aged (45-day-old at 29°C) flies. A marked reduction in Med19 protein levels is observed in aged flies compared to tubulin controls. (A’) Quantification of Med19 protein levels from the Western blot. The box plot shows a significant decrease in Med19 expression in older flies compared to younger ones (p < 0.05), indicating age-dependent downregulation of *Med19*. (B) Schematic of vitality tests performed at different ages, comparing 80 flies expressing *UASMed19* upon *ubiquitin* promoter versus *UASlacZ* controls. (B’) Med19 upregulation significantly increases the vitality of aged flies. Negative geotaxis tests show that Med19 overexpression improves climbing efficiency at 25 days old (*** Unpaired t-test: P-value <0,05) and 30 days old (* Unpaired t-test: P-value <0,05) compared to controls. (C) Schematic of the experimental setup in which Med19 is overexpressed in frail flies using the *UASMed19* construct. Flies were raised at 18°C and shifted to 29°C on day 62 to induce *Med19* upregulation. (C’) Survival analysis shows that Med19 overexpression (red line) improves survival compared to control flies (black line). Median lifespan during the induction period increased from 16 days to 19 days in controls (****P-value), demonstrating that boosting Med19 levels can partially rescue age-related frailty and extend lifespan.

Given that frailty is associated with a decline in vitality, muscle function (sarcopenia), and locomotion, we asked if maintaining high levels of Med19 expression could protected against this deterioration of individual fitness. One way to measure this loss of vitality in *Drosophila* is the negative geotaxis test (equivalent of the grip strength test in humans). As flies age, their ability to climb decreases significantly, reflecting reduced performances and overall frailty, hallmarks of “unfit aging”. We therefore performed negative geotaxis tests on Med19 overexpressing flies and found that climbing activity was unchanged in young (7 days-old) flies. However, after 25 days of life at 29°C, at the transition phase between healthy aging and disability, almost 70% of Med19 overexpressing flies was able to climb up to the tube compared to around 50% for the control population (Figure 6B’). This difference was also observed at 30 days of life at 29°C when age-related decline becomes more apparent, where almost 40% of Med19 overexpressing flies climbed compared to 20% of control flies. These findings indicate that Med19 protein level decreases with age and that genetically counteracting this decline preserves muscle capacity supporting the role of *Med19* in healthy aging.

### Maintaining Med19 levels extends survival in frail individuals

In *Drosophila*, the onset of frailty—representing a transitional phase between healthy aging and disability—remains poorly documented. Nevertheless, significant physical deterioration has been reported between 30 and 45 days of life, along with a marked increase in mortality rate (“break day”) beginning around day 30 at 25 °C (Shahrestani et al. 2016; Rhodenizer et al. 2008; Rose et al. 2002; Grotewiel et al. 2005).

To assess whether Med19 upregulation can counteract age-associated decline, we performed rescue experiments at an advanced age, when physical deterioration begins to manifest. As shown in Figure 6C-C’ and S1 table, we upregulated Med19 in “frail” individuals (60 days-old at 18°C - equivalent to ∼30 days at 25°C) and showed that Med19 upregulation significantly improves median lifespan by 19% from the induction time at 29°C (+19 days with *ubMed19* instead of 16 days for the control – p < 0,0001). As previously observed, Med19 overexpression did not modify the rate of mortality once it began to rise, but shifted the inflection point to a later time, indicating a delayed onset of frailty (Figure 6C’ and S1 table). These results indicate that restoring Med19 levels, even at advanced ages, can mitigate age-related functional decline during the early phase of frailty. They further suggest that the physiological decline of *Med19* expression could be a causal factor in the transition from healthy aging to disability.

## DISCUSSION

### Med19, a frailty rheostat regulating organismal and cellular fitness

Aging in *Drosophila melanogaster*, like in vertebrates, is characterized by a gradual decline in physiological functions culminating in a critical “frailty point,” at which survival rates dramatically worsen. In this work, we demonstrate for the first time a pro-longevity action of the Mediator complex subunit Med19. Constitutive upregulation of Med19 in *Drosophila melanogaster* nearly doubles median lifespan and delays the onset of frailty, thus positioning Med19 alongside canonical modulators of lifespan such as Indy, FOXO, JNK or the Insulin/IGF-1 pathway (Rogina et al. 2000; Wang et al. 2003; Wang et al. 2005; Tatar et al. 2001). Moreover, the observed improvement in vitality shows that Med19 upregulation not only enhances lifespan but also healthspan. By contrast, Med19 knockdown drastically reduces lifespan and results in an earlier acceleration of mortality. Beyond interventions that merely delay the onset of specific pathologies, Med19 appears to strengthen fundamental cellular defenses, particularly under DNA-damaging or oxidative conditions. This mediator subunit is thus closely associated with the preservation of genomic stability and stress resistance — processes whose decline is a hallmark of aging.(López-Otín et al. 2013). Importantly, our observation that Med19 protein levels naturally decline in older flies suggests that this late-life drop could be one cause of age-related frailty.

At the cellular level, Med19-depleted cells behave as “losers” in cell competition assays, undergoing azot- and JNK-dependent apoptosis upon contact with wild-type neighbors. Although viable in a weakened environment (e.g., *Minute* background), Med19-depleted cellular clones are outcompeted in a wild-type context, underscoring the role of Med19 in sustaining cell fitness. This supports the notion of “cellular frailty,” where compromised cells are selectively eliminated to maintain tissue homeostasis, echoing the frailty syndrome at the organismal level.

### Med19 as an evolutionarily-conserved cellular stress protector

Interestingly, while loss of Med19 enhances the transcription of stress-response pathways (e.g., iron–sulfur cluster biogenesis, glutathione metabolism, and DNA damage response), it weakens stress resistance at the whole-organism level. One hypothesis that may account for this apparent paradox is that knocking down Med19 elevates intrinsic stress leading to compensatory but yet insufficient transcriptional changes that ultimately fail to protect the organism. This scenario aligns with our observation that at the opposite, Med19 overexpression strongly protects flies from stress, likely by preventing the accumulation of reactive oxygen species (ROS) and DNA damage. This interpretation is also consistent with the results of our cell competition assays in which Med19 downregulation induces an Azot-dependent “loser” phenotype, known to display chronic activation of stress response pathways including the oxidative stress response and the JNK pathway, a key regulator of stress signaling (Kucinski et al. 2017; Kodra et al. 2024; Igaki 2009).

How does Med19 depletion trigger cellular stress? One possible mechanism arises from a recent study by Perrimon’s group, which showed that Wingless represses genotoxic DNA damage in *Drosophila* wing discs (Ewen-Campen & Perrimon 2024). In line with this, we recently found that Med19 knock-down leads to a loss of Wingless expression in these tissues, potentially resulting in genomic instability (Jullien et al. 2022). Alternatively, an increase in the expression of genes related to Fe**-**Sulfur cluster biogenesis may indicate that cells are attempting to compensate for the loss or dysfunction of these clusters. Such dysfunction could compromise multiple enzymes essential for mitochondrial respiration, DNA repair, and redox homeostasis, leading to reduced energy production and genomic instability. As a result, cells accumulate higher levels of reactive oxygen species and experience molecular damage, causing severe cellular stress (Rouault & Maio 2017).

Multiple lines of evidence suggest that Med19’s role as a stress modulator may be evolutionarily conserved. In yeast, the Med19 homolog Rox3 was shown to be recruited to stress-regulated promoters and to function in both heat shock and osmotic stress tolerance (Evangelista et al. 1996; Singh et al. 2006; Gulshan et al. 2005). In plants, Med19A responds to nitrogen deficiency through phase separation, coordinating senescence-specific gene regulation (Cheng et al. 2022). In humans, Med19 is upregulated in several cancers (Zhang et al. 2021; Jin et al. 2024). Given that genomic instability and oxidative stress are hallmarks of cancer (Negrini et al. 2010; Hanahan & Weinberg 2011), our findings in *Drosophila*— demonstrating Med19’s role in protecting against DNA damage and oxidative stress—suggest that its upregulation in tumors may reflect a conserved function that enables cancer cells to cope with the chronic stress conditions characteristic of malignant progression.

### Mediator subunit specificity

Med19 functions through the Mediator complex, which consists of 30 distinct subunits. This raises the question of whether its role in cellular and organismal frailty reflects a general function of the complex in regulating RNAPolII-dependent transcription or a more specific function of Med19. Although nearly all subunits are always present within the complex, they are not functionally equivalent. For example, while loss of Med14 broadly downregulates RNAPolII activity, our data show that only 9% of genes are misregulated either positively or negatively in Med19-KO wing pouch (El Khattabi et al. 2019). Similarly, mutant clones for other MED subunits display distinct cellular phenotypes: Med17 loss induces cell death in all tissues, whereas loss of either Med12, Med13, Cdk8 or CycC does allow normal cell proliferation (Boube et al. 2000; Loncle et al. 2007; Treisman 2001). The tissue-specific lethality seen with *Med19* mutants suggests a unique role in cell competition. Of note, several other Mediator subunits are implicated in response to stress in either nematodes, plants, or mice (Goh et al. 2014; Dard et al. 2024; Chen et al. 2022). Whether some of these subunits act together with Med19 in stress-induced transcriptional regulation in *Drosophila* and other species is one next step to understand.

### Med19’s expression dynamics and frailty

While the Mediator complex subunits are often considered ubiquitously expressed, transcriptomic data from the Fly modENCODE and Fly Cell Atlas projects reveal subtle changes in *Med19* mRNA or Med19 protein across tissues and developmental stages, and proteomic analyses further indicate that Med19 levels may shift under stress conditions (FlyBase Gene Report: Dmel\MED19; (Chen et al. 2022; Kolonay et al. 2024)). We detect a significant Med19 decline in very old *Drosophila melanogaster* adults, which correlates with heightened frailty, yet adult-specific reintroduction of Med19 in already aged flies can still partially restore longevity. These results highlight that post-developmental role of Med19 are highly relevant to healthy aging and raise questions about whether comparable shifts in Med19 expression occur in mammalian tissues. A deeper understanding of Med19 regulation—both transcriptional and post-translational—will be crucial for clarifying how this subunit contributes to tissue homeostasis and for exploring its therapeutic potential.

Overall, this work broadens our understanding of Mediator subunits beyond general gene regulation, demonstrating that Med19 is central to stress defense, genomic stability, and longevity in *Drosophila*. Deciphering the upstream signals and downstream effectors of Med19, in tandem with investigating possible roles in human aging, should further illuminate how this key transcriptional cofactor intersects with frailty and healthspan.

## MATERIALS AND METHODS

### *Drosophila* stocks and reagents

#### KEY RESOURCES TABLE

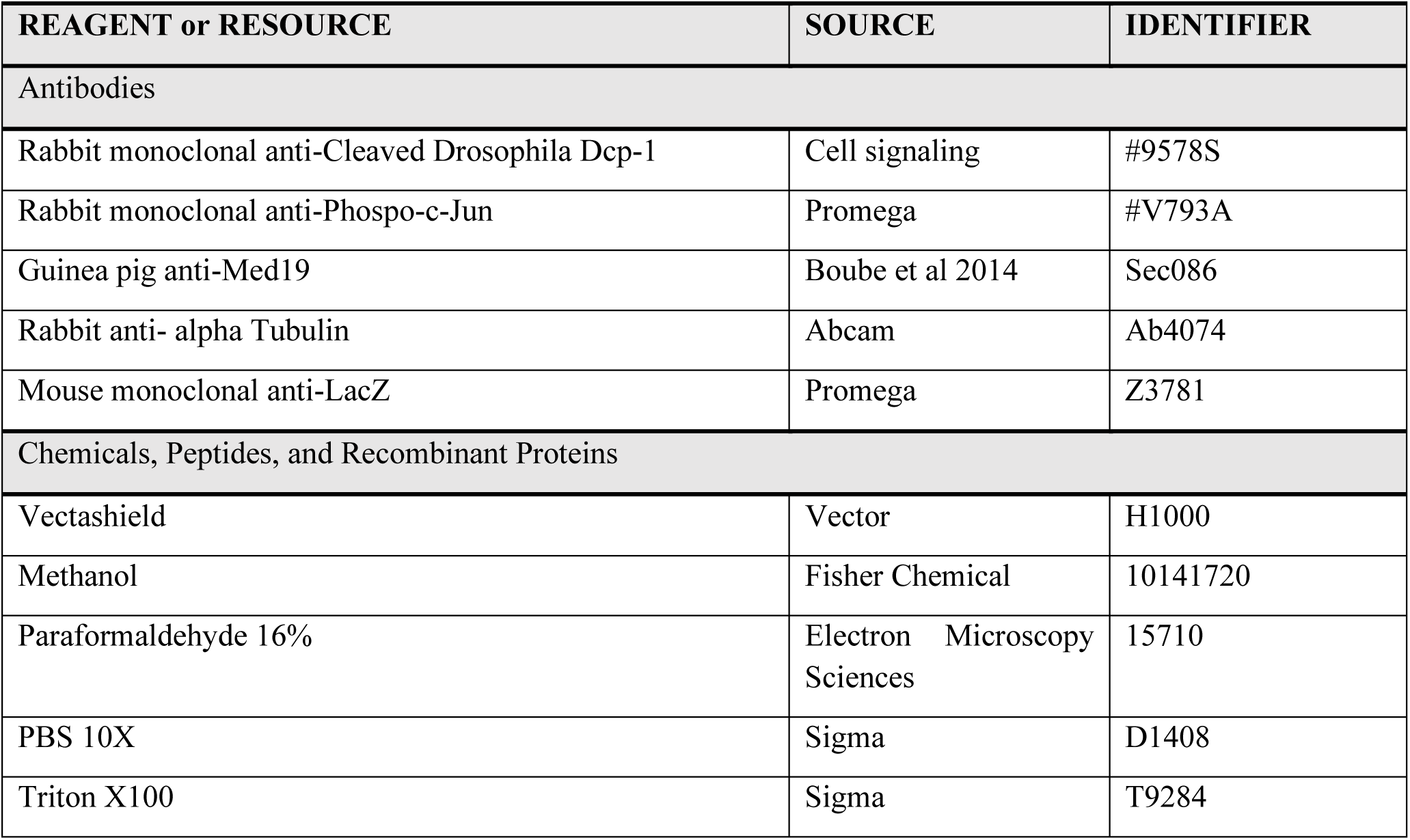
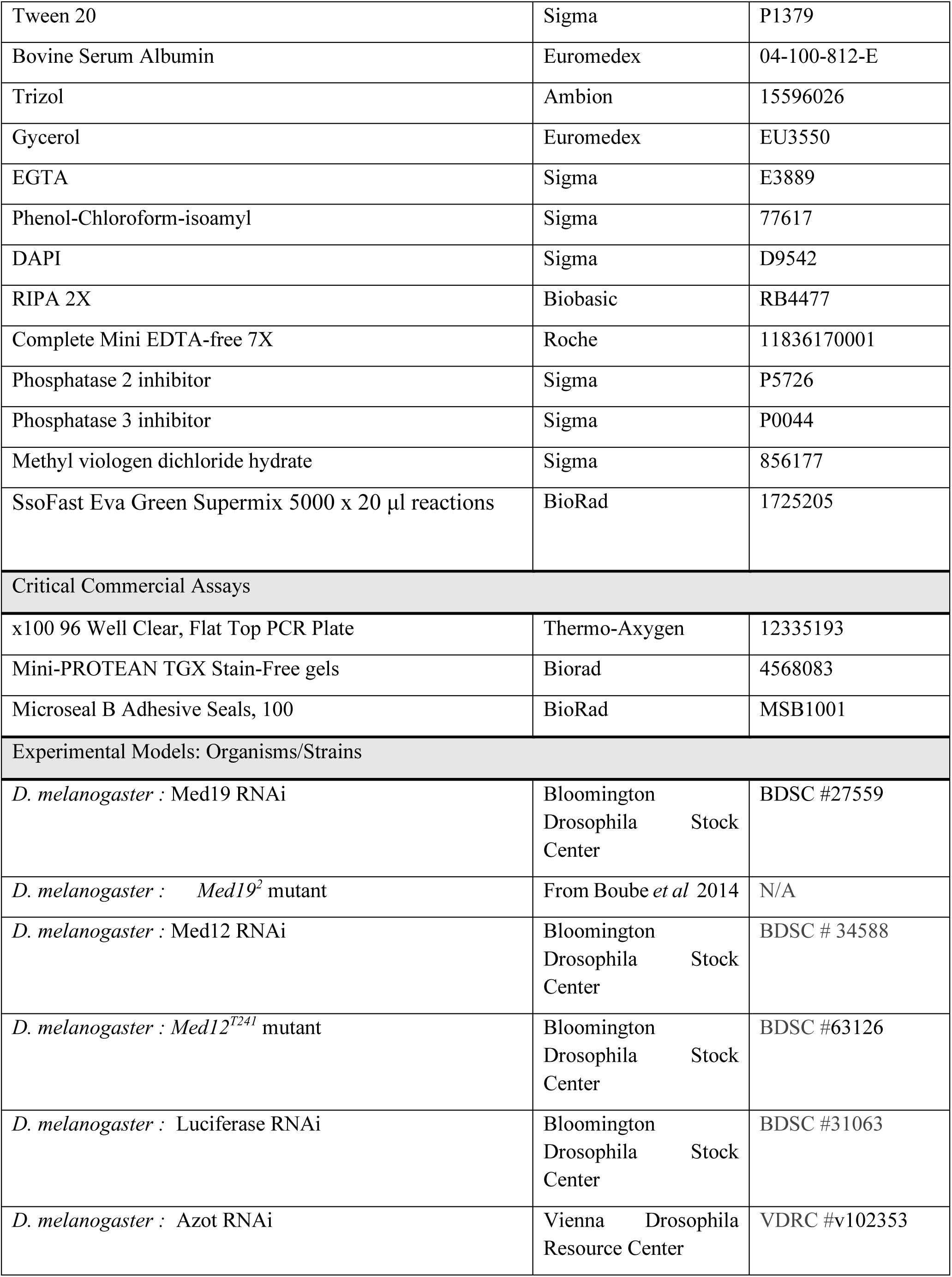
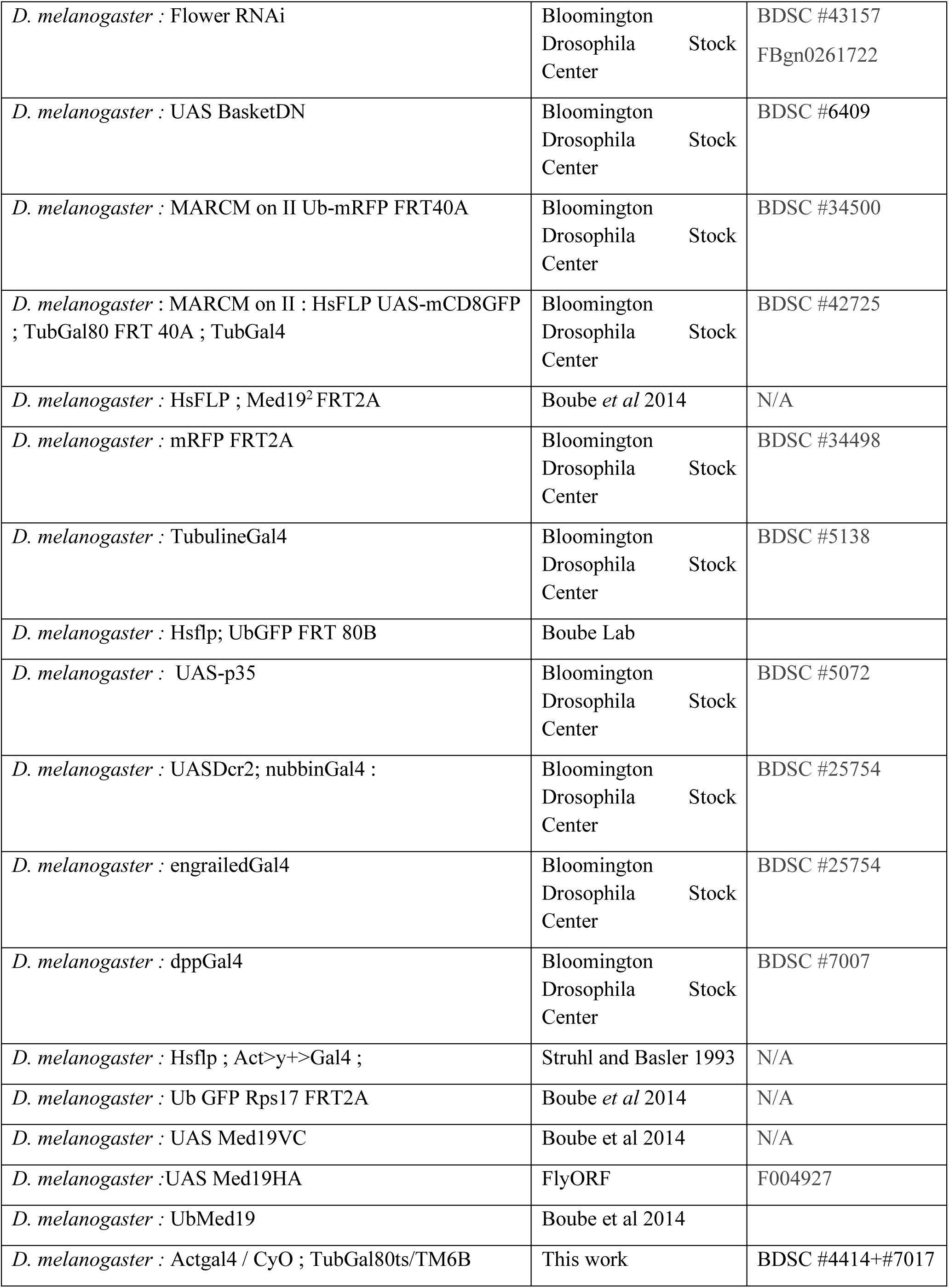
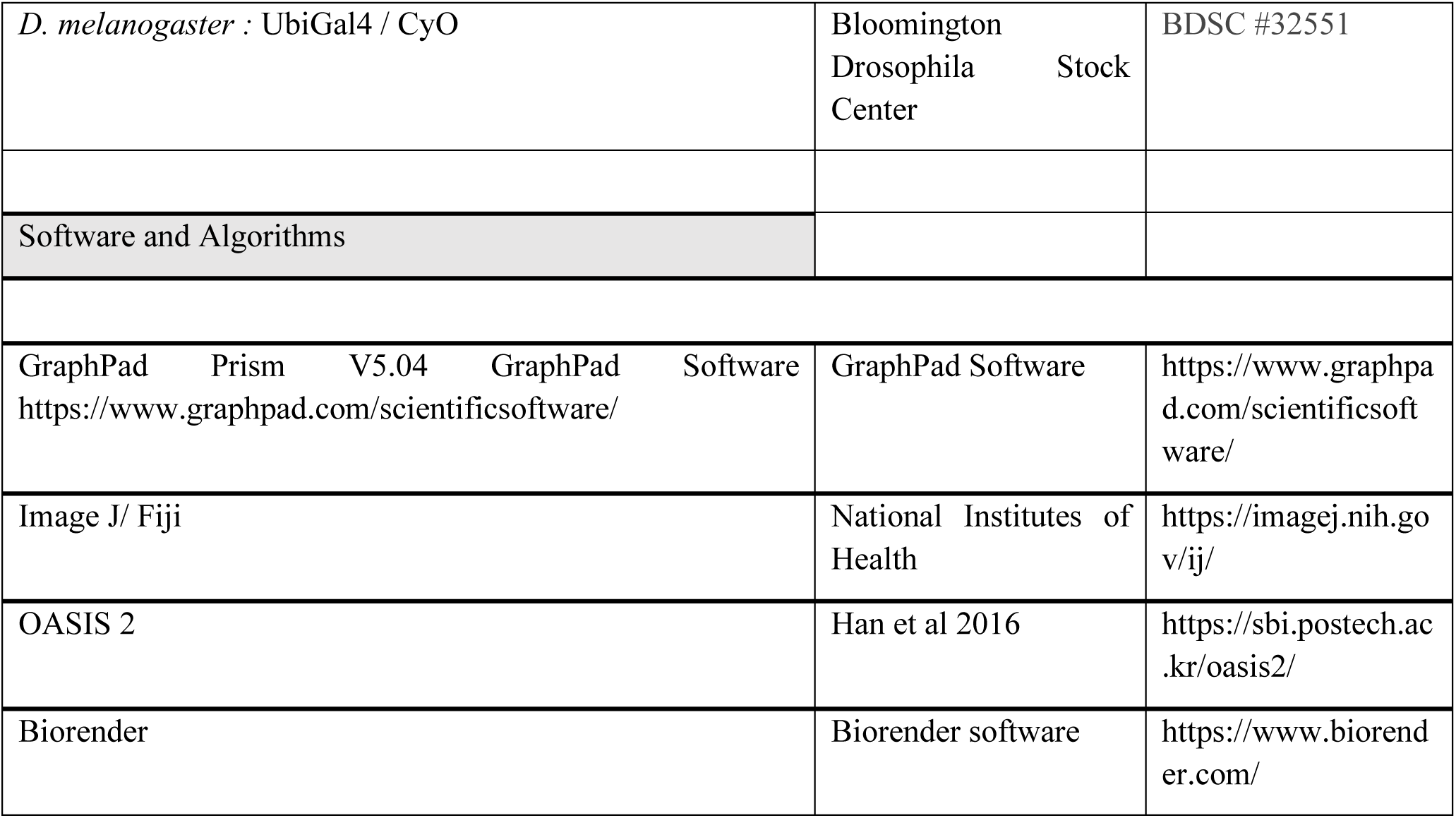

### MARCM Experiments

Females P{hsFLP}1, y1 w* P{UAS-mCD8::GFP.L}Ptp4ELL4; P{tubP-GAL80}LL10 P{neoFRT}40A; P{tubP-GAL4}LL7 is crossed with males from y[1] w[1118]; P{w[+mC]=Ubi-mRFP.nls}2L P{ry[+t7.2]=neoFRT}40A/CyO; UAS Med19 RNAi or y[1] w[1118]; P{w[+mC]=Ubi-mRFP.nls}2L P{ry[+t7.2]=neoFRT}40A/CyO; UAS-Luciferase RNAi. Heat-shocks are performed 48H after egg laying during 10’ at 37,5°. After heat-shock, flies are raised at 29°C in order to enhance RNAi expression. Fixation and dissection are performed at least 48H hours after the heat-shock.

### Mitotic clones

Mitotic clones were induced by Flp recombinase expressed from a *hsp70-Flp* transgene on heat induction in L1 and L2 1 hour each at 37,5°. Fixation and dissection in the late L3 stage.

Mitotic clones were also induced by Flp recombinase expressed from a UAS-Flp element under Gal4 control as indicated above (en.Flp; ap.Flp). Clones were generated and identified in marked progeny from crosses using the following stocks:

### Immunostaining

Immunostaining was performed using the following antibodies: anti-activatedDcp1 (1:200), antiMed19 (1:1000), anti-LacZ (1:1000), anti ^P^Jun (1:500). Larvae were collected and fixed for 20min in PAF 4%, PBS 1X, EGTA. Following fixation larvae were rinsed at least three times in methanol and stored a -20°C in fresh methanol until required. Fixed larvae are rehydrated in 75%, 50% and 25% methanol diluted in PBS 1X, Triton 0,1% (PBST). Fixed larvae were then blocked in PBS with 0,1% triton-X detergent and 1% bovine serum albumin (PBST-BSA) for 3x20min. Larvae were then incubated with diluted primary antibody (at appropriate concentrations) in PBST-BSA overnight at room temperature or 48h at 4°C. After that, the primary antibody solution was removed and larvae washed three times in PBST-BSA for a total of 1h before incubation with the appropriate secondary antibody (diluted at 1:500 in PBST-BSA) for 2h at room temperature. Following secondary antibody staining, larvae are rinsed in PBS, Tween 0,3% solution and stored at 4°C in fresh PBS-Tween or PBS-Glycerol 50% for a maximum of 7days. Dissection and mounting of imaginal discs were performed in PBS-Glycerol or Vectashiel solution and imaging was performed on Leica SP5 or SP8 microscopes.

### RNAseq

We choose RNAi-induced depletion with a strong *nubGal4* driver line and a *UAS-DRC2* transgene potentializing RNAi activity in larval imaginal discs. This strong Med19 depletion allows to detect the regulatory activity of the Med19 subunit in a tissue that cannot develop until adult stage in these conditions. Female L3 larvae were hand-dissected in PBS 1X Tween 0.1 % at 4°C to recover wing imaginal discs. Each disc was cut to discard the notum part and keep the wing pouch. A mean of 75 wing pouches were used per sample: *nubGAL4 UAS-DCR2 UAS_Med19 RNAi* vs control *nubGAL4 UAS-DCR2 UAS-LucRNAi*. Each sample was replicated in 3 independent experiments. RNA was extracted from discs using TRIzol reagent, selected for polyadenylation and paired end sequenced on a NovaSeq 6000 (Illumina Genewiz). The bioinformatic analysis was performed using our locally installed Galaxy instance (http://sigenae-workbench.toulouse.inra.fr). First, the quality control of sequences was done with FastQC (v0.11.4), then reads were aligned using TopHat2 (v2.0) on the *Drosophila melanogaster* genome dm6. Paired reads were counted by gene using htseq (v1.0). These counts were used to perform gene-level differential expression analyses using DESeq2 (1.26.0) in R (v3.6.1) between *nubGAL4 UAS-Med19* RNAi vs *nubGAL4 UAS-Luciferase RNAi*. We selected differentially expressed genes (DEGs) by setting first a significance determined by Benjamini-Hochberg adjusted P-values < 0.05 and second a log2 fold change of their expression > 0.5 or < - 0.5.

Functional annotation of DEGs was done by using Gprofiler (https://biit.cs.ut.ee/gprofiler/gost) to find enriched GO terms belonging to Biological processes, and to generate GO terms tree. List of genes annotated in the GO: 0006950 “Cellular response to stress” in Gprofiler and Flybase have been compared to our DEGs upregulated genes. Heatmaps have been generated using R software.

### Western Blot

Flies were cultured at 29°C degrees in the incubator, and transferred to a fresh medium every 3 days. To compare Med19 expression of young and old age fruit-flies, we collected young adults at five days post eclosion and old flies at day 45 days post eclosion. In each experiment 12 to 20 flies (50% males and 50% females) were dissected to keep head and thorax only, samples were fast frozen using dry ice and kept to -80°C until protein extraction. Protein extraction was performed using RIPA 2X and gel migrations using Biorad TGX Stain-Free Precast gels (4-20% gradient gel). Staining was performed using antiMed19 (1:500) or antiTubulin (1:5000) diluted in BSA 1%. Med19 protein levels were quantified from three independent experiments normalized to tubulin levels.

### Lifespan experiment

Flies were reared on standard food medium (sugar (40g/l), 2% yeast (28g/l), cornmeal (74g/l), agar(7,5g/l), Moldex (2g/l), propionic acid (4ml/l). Experiments were performed at 25°C for *ubMed19*, or at 18°C and then shifted at 29°C for adult specific expression of *Med19* cDNA or RNAi. Given that *Drosophila* lifespan is sensitive to temperature (maximum lifespan ∼120 - 140 days at 18°C, ∼80 days at 25°C and ∼40-50 days at 29°C Molon 2020), we only compared data sets within the same experiment. For lifespan experiments at 25°C, 3 to 5 tubes of 20 to 30 individuals (half males and half females) were used per genotype (see S1 table, for details) and each experiment were repeated 2 or 3 times. Flies were then transferred to a fresh medium every 2 or 3 days and dead flies were counted at this time. Kaplan -Meier survival curves were generated with GrafPad Prism software and in-depth statistical analyses of lifespan curves with OASIS2 software (https://sbi.postech.ac.kr/oasis2/). For each experiment, table entitled “Mean/median lifespan” indicates the number of subjects, the mean lifespan (green rectangle) with standard deviation and 95% Confidence Interval. Median lifespan, corresponding to the day when 50% mortality passed is framed in blue, and maximum lifespan at 90% mortality, in magenta. Log rank tests were used to compare entire survival curves and mean lifespan (dashed green) whereas Mann and Witney U tests were applied to compare median (dashed blue) and maximum (dashed magenta) lifespan at 90% mortality. P-value > 0.05: ns, P-value < 0.01: **, P-value < 0.001: ***, P-value < 0.0001: ****.

### Negative geotaxis tests

To assess the locomotor activity of *Drosophila*, we used negative geotaxis tests measuring fly wall-climbing reflex after a fall. The test measures the number of flies that successfully climb, without flying, half the height of the tube (ie >4.5 cm) within 5 seconds. Experiments were performed at 18°C and then shifted at 29°C for adult specific expression of Med19 cDNA. For each genotype (UAS lacZ control vs UAS Med19), we use 4 tubes of 20 flies (10 males and 10 females), and the test is performed 3 times on each tube to ensure representativity. The test is carried out at 10, 25, and 30 days of fly life at 29°C, starting with 20 flies per tube. The analysis of the videos was conducted retrospectively one by one, using Windows Media Player software following single individual to avoid inter-individual analysis biases. The results were processed using Excel software to calculate the percentage of flies reaching half the height of the tube.

### Paraquat exposure

We used 20 mM Paraquat (Methyl viologen dichloride hydrate) diluted in 5% sucrose solution. Flies (3 tubes of 10 males + 10 females) were placed in an empty tube with a bench coat humidified with the paraquat solution for 24 hours (Figure 2B, D). Flies were placed into fresh medium and then transferred and counted every 2 to 3 days.

### UV Irradiation

UV irradiation was performed using a Stratalinker UV Crooslinker 1800 (Stratagene). We exposed 3 tubes of 10 males + 10 females each to 48 x 10^2^ Joules/m2 (Figure2A, C). Flies were placed into fresh medium and then transferred and counted every 2 to 3 days.

## ACKNOLEDGEMENTS

We are grateful to M. André for technical help in Western experiments, J. Favier, A. Destenabes, V. Nicolas and M. Crozatier for CBI Fly facilities, and M. André, A.Vincent, F. Payre, M. Crozatier, and C. Monod for critical comments and valuable help. We thank Toulouse RIO imaging platform for assistance with confocal microscopy. We are grateful to the genotoul bioinformatics platform Toulouse Midi-Pyrenees and Sigenae group for providing help and storage resources on the local Galaxy instance http://sigenae-workbench.toulouse.inra.fr. We acknowledge the big-A facility of CBI that helped us a lot in bioinformatic and biostatistical analysis. We deeply thank Eduardo Moreno, Gines Morata, Dirk Bohmann, the Bloomington *Drosophila* Stock Center and the Developmental Studies Hybridoma Bank for providing us with antibodies, plasmids and fly stocks.

## CRediT author contributions

Conceptualization AP, CS, MB

Methodology AP, SBF, EG, MB

Software

Data curation EG

Investigation AP, EG, SBF, CDB, DJ, MB

Validation AP, EG, SBF, CDB, DJ, MB

Formal analysis AP, EG, XC, AR, MB

Supervision CS, HMB, MB

Funding acquisition CS, HMB, MB

Visualization AP, MB

Project administration MB

Resources CS, HMB, MB

Writing – original draft AP, MB

Writing – review & editing AP, EG, DJ, JLF, CS, MB

## Supporting information legends

**S1 Table - OASIS2 statistical analysis of lifespan curves.**

For each experiment, the table entitled “Mean/median lifespan” indicates the number of subjects, the mean lifespan (green rectangle) with standard deviation and 95% Confidence Interval. Median lifespan, corresponding to the day when 50% mortality passed is indicated by blue rectangles, and maximum lifespan at 90% mortality by magenta rectangle. Dashed green rectangles indicates Log rank statistical tests to compare mean lifespan data, dashed blue and magenta rectangles indicates statistical results from Mann and Witney U tests to compare median and maximal 90% lifespan, respectively. *P-value > 0.05: ns, P-value* < 0.01: **, *P-value* < 0.001: ***, *P-value* < 0.0001: ****). Maximum lifespan could be determined by "fundamental process of aging" whereas mean lifespan changes with various condition such as diseases. Thus, increasing maximal lifespan may be an indicator that an intervention is slowing the general processes of aging and not merely retarding the development of specific diseases.

**S2 Table - Med19 target genes involved in DNA Damage Response and sulfur compound metabolic process**

**S3 Figure.**
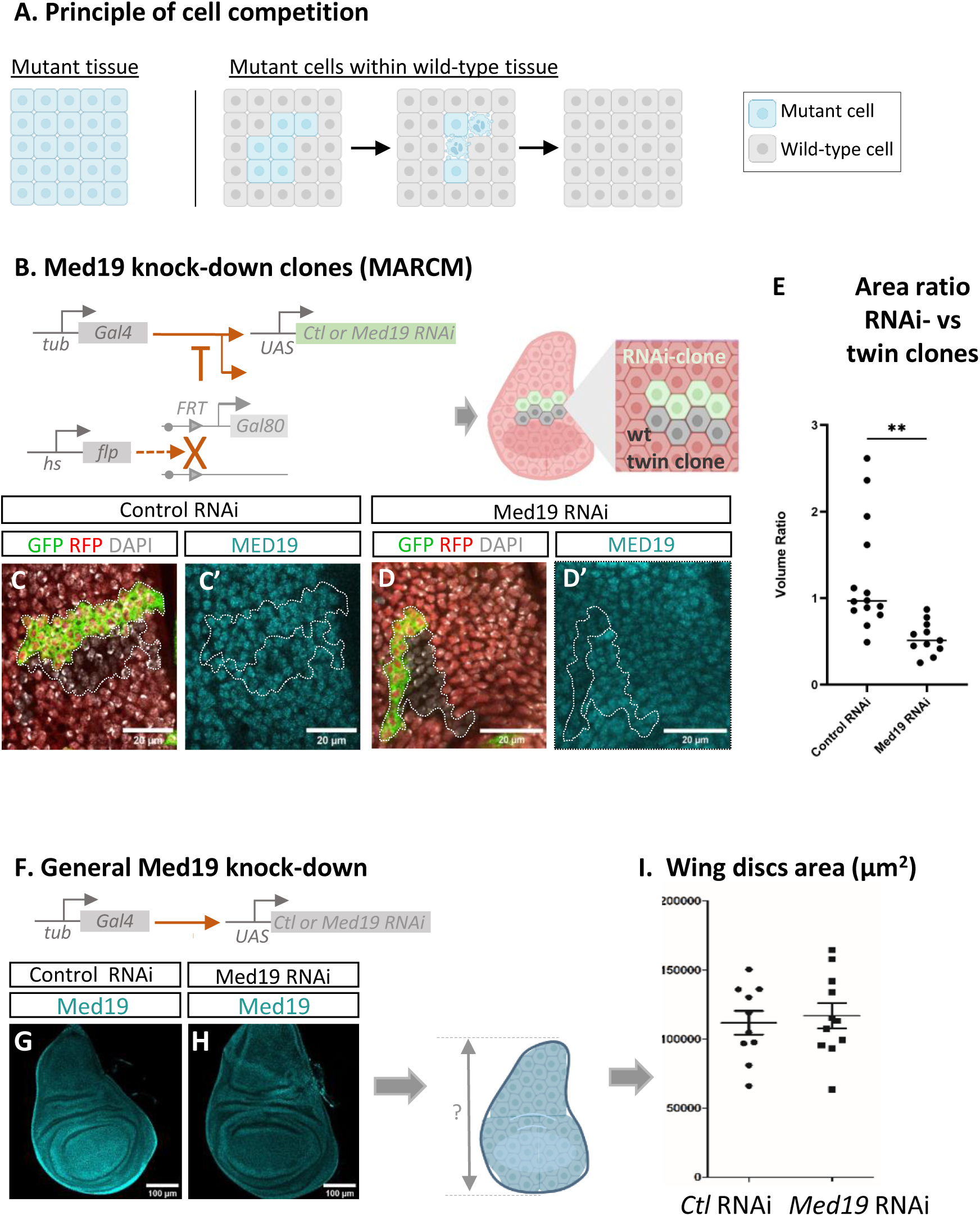
At the cellular level, Med19-depleted cells display a loser cell phenotype. (A) General principle of cell competition, a natural process allowing active elimination of unfit “loser cells” by neighboring, more ‘fit’ or ‘winner’ cells. Whereas unfit mutant cells can proliferate normally when surrounded by cells of similar status (left panel), they behave as “losers” and are eliminated through cell competition when surrounded by healthy wild-type neighbors (right panels). (B-E) Med19 knockdown clones (MARCM System). (B) Schematic of the MARCM system. MARCM clones expressing RNAi against control (C-C’) or Med19 (D-D’), together with *UAS-GFP* under the *tub-GAL4* control. Twin wild-type clones (RFP-) and heterozygous surrounding cells (RFP+) bear a *Ub-Gal80* transgene inhibiting GAL4 expression in these cells. Nuclei are marked with DAPI staining (grey). Panels C’ and D’ shows Med19 immunostaining (E) Quantification of ratio between Luc RNAi- or Med19 RNAi-expressing twin clone areas. With control RNAi (C-C’), GFP+ and GFP-clones display similar size whereas in (D-D’) Med19 knock-down clones (GFP+ in D and weaker Med19 signal in D’) appears smaller than their corresponding wild type clone (RFP – GFP - in D and stronger Med19 signal in D’). (F-I) Med19 depletion in whole imaginal discs. (F) Schematic of the experimental design for global Med19 knock-down using the *tub-Gal4* > *UAS-Med19* RNAi, leading to uniform depletion of Med19 throughout the wing disc. (G-H) Confocal images show that wing discs with global Med19 RNAi expression (G) are similar in size to control discs (F), despite the reduced Med19 signal. (I) Quantification of wing disc area reveals no significant difference between control and Med19 RNAi discs, suggesting that Med19-depleted cells survive and proliferate normally when neighboring cells are similarly depleted, contrasting with the selective elimination seen in mosaic clones

**S4 Figure.**
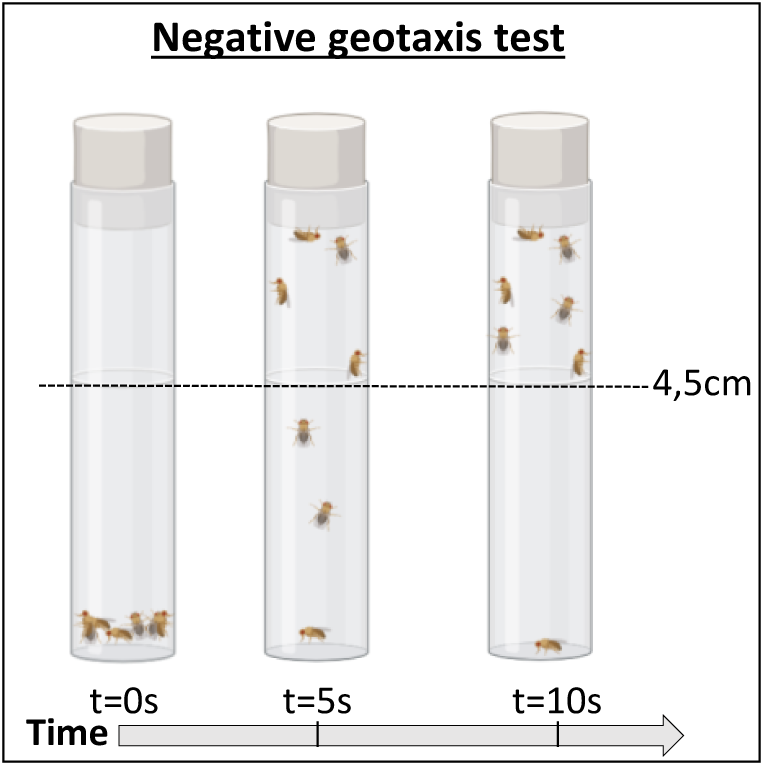
Schematic of the experimental design for negative geotaxis tests.

